## Supplementary Information for "The neurocomputational link between defensive cardiac states and approach-avoidance arbitration under threat"

**Supplementary Table S1. Coordinates and statistics of positive and negative correlations between BOLD and reward/threat magnitudes during the anticipation screen**

| Region | Side | x, mm | y, mm | z, mm | Cluster size, $mm^3$ | Peak T | *p* value |
|---|---|---|---|---|---|---|---|
| *Reward – positive* | | | | | | | |
| Occipital cortex | L/R | -16 | -102 | -2 | 58,808 | - | < .001 |
| Amygdala | L | -20 | -2 | -14 | 1,672 | - | = .003 |
| Precentral gyrus | L | -36 | -22 | 60 | 1,576 | - | < .001 |
| Supplemental motor area | L/R | 2 | -8 | 58 | 4,048 | - | < .001 |
| Ventral striatum | L | -6 | 10 | -4 | - | 4.57 | = .003* |
| | L | -16 | 10 | -12 | - | 3.60 | = .050* |
| | R | 8 | 8 | -2 | - | 5.26 | < .001* |
| | R | 10 | 12 | -10 | - | 4.47 | = .004* |
| vmPFC | R | 4 | 64 | -2 | - | 4.16 | = .059* |
| *Reward – negative* | | | | | | | |
| Lateral orbitofrontal cortex | R | 38 | 42 | -10 | 1,120 | - | = .024 |
| *Threat – positive* | | | | | | | |
| Occipital cortex | L/R | -18 | -78 | -12 | 30,712 | - | < .001 |
| SMA/dACC | L/R | 10 | 10 | 46 | 11,872 | - | < .001 |
| | L | -2 | 2 | 44 | - | 4.68 | = .025* |
| | R | 10 | 10 | 44 | - | 5.99 | < .001* |
| | R | 8 | 26 | 30 | - | 4.95 | = .011* |
| | R | 14 | 20 | 32 | - | 4.79 | = .018* |
| | R | 10 | 24 | 34 | - | 4.76 | = .020* |
| Insular cortex | L | -42 | 16 | 2 | 3,512 | - | < .001 |
| | R | 34 | 18 | -8 | 7,000 | - | < .001 |
| Postcentral gyrus | L | -44 | -26 | 52 | 4,744 | - | < .001 |
| dlPFC | R | 34 | 32 | 36 | 5,568 | - | < .001 |
| | L | -32 | 46 | 24 | 2,240 | - | < .001 |
| Precuneus | L/R | 6 | -54 | 54 | 6,528 | - | < .001 |
| Temporoparietal junction | L | -58 | -48 | 42 | 3,032 | - | < .001 |
| | R | 62 | -44 | 32 | 4,504 | - | < .001 |
| Intraparietal sulcus | L | -18 | -74 | 48 | 1,240 | - | < .001 |
| *Threat – negative* | | | | | | | |
| vmPFC | L/R | -4 | 50 | -10 | 2,016 | - | < .001 |
| *Rew-by-Thr – negative* | | | | | | | |
| Occipital cortex | L/R | -6 | -80 | -6 | 3,464 | - | < .001 |

*All coordinates are defined in MNI152 space. All listed statistics are significant at p < .05 FWE-corrected at the cluster-level (whole-brain) or peak-level small volume corrected (FWE-SVC, predefined ROIs, indicated with asterisks '*'). dACC: dorsal anterior cingulate cortex; SMA: supplemental motor area; dlPFC: dorsolateral prefrontal cortex; vmPFC: ventromedial prefrontal cortex.*

**Supplementary Table S2. Coordinates and statistics of (de)activations as a function of passive and active approach vs. avoidance choices during the anticipation screen**

| Region | Side | x, mm | y, mm | z, mm | Cluster size, mm$^3$ | Peak T | *p* value |
|---|---|---|---|---|---|---|---|
| *Approach > Avoid* | | | | | | | |
| Occipital cortex | L | -20 | -98 | 2 | 6,216 | - | < .001 |
| | R | 28 | -98 | -6 | 2,208 | - | < .001 |
| Hippocampus/PHG/AMY | R | 22 | -10 | -14 | 16,192 | - | < .001 |
| Amygdala | L | -18 | -2 | -22 | - | 4.37 | = .005* |
| | L | -28 | -6 | -12 | - | 4.09 | = .011* |
| | L | -16 | -4 | -18 | - | 3.98 | = .015* |
| | L | -16 | -2 | -14 | - | 3.95 | = .016* |
| | L | -24 | 0 | -12 | - | 3.79 | = .024* |
| | L | -12 | -2 | -16 | - | 3.53 | = .048* |
| | R | 24 | -8 | -14 | - | 5.44 | < .001* |
| | R | 30 | -4 | -18 | - | 4.82 | = .001* |
| | R | 20 | -2 | -18 | - | 4.14 | = .009* |
| | R | 26 | 4 | -28 | - | 3.78 | = .025* |
| | R | 32 | 4 | -26 | - | 3.27 | = .090* |
| Precentral gyrus | L | -32 | 24 | 62 | 6,384 | - | < .001 |
| | L | -54 | 2 | 40 | 992 | - | = .036 |
| | R | 56 | 6 | 40 | 960 | - | = .041 |
| Supplemental motor area | L/R | 2 | -16 | 64 | 8,544 | - | < .001 |
| Parietal operculum | R | 56 | 2 | 6 | 944 | - | = .044 |
| vmPFC | L/R | -4 | 64 | -6 | 5,504 | - | < .001 |
| Ventral striatum | L | -6 | 8 | -4 | - | 5.72 | < .001* |
| | L | -8 | 8 | -10 | - | 4.85 | = .001* |
| | R | 8 | 8 | -4 | - | 5.54 | < .001* |
| | R | 6 | 12 | -4 | - | 5.54 | < .001* |
| BNST | L | -6 | 4 | -4 | - | 3.62 | = .010* |
| | L | -4 | 0 | 0 | - | 3.08 | = .040* |
| | R | 8 | 4 | -4 | - | 4.05 | = .003* |
| | R | 6 | 0 | -2 | - | 3.71 | = .008* |
| *Approach < Avoid* | | | | | | | |
| Precuneus | R | 6 | -56 | 54 | 3,520 | - | <.001 |
| *Active > Passive* | | | | | | | |
| Postcentral gyrus | L | -34 | -22 | 44 | 4,728 | - | < .001 |
| Thalamus | L/R | 4 | -18 | 0 | 872 | - | = .030 |
| $Choice_{Passive} > Choice_{Active}$ | | | | | | | |
| Lingual gyrus | L/R | -2 | -66 | 6 | 960 | - | = .021 |
| Supplemental motor area | L/R | -4 | -4 | 64 | 4,416 | - | < .001 |
| Postcentral gyrus | L | -40 | -22 | 56 | 1,432 | - | = .002 |
| $Choice_{Passive} < Choice_{Active}$ | | | | | | | |
| Occipital cortex | L | -16 | -92 | -8 | 2,016 | - | < .001 |
| | R | 20 | -90 | -4 | 2,464 | - | < .001 |

*All coordinates are defined in MNI152 space. All listed statistics are significant at p < .05 FWE-corrected at the cluster-level (whole-brain) or peak-level small volume corrected (FWE-SVC; predefined ROIs, indicated with asterisks '*'). PHG: parahippocampal gyrus; AMY: amygdala; vmPFC: ventromedial prefrontal cortex; ACC: anterior cingulate cortex; BNST: bed nucleus of the stria terminalis.*

*Similar neural effects for anticipation of passive vs. active approach-avoidance choices*

We explored whether the observed neural approach-avoidance circuit (see main text) was differentially involved in passive vs. active approach-avoidance choices. The choice-by-response interaction indicated a significant effect in the SMA ($p$ < .001 cluster-level FWE), left postcentral gyrus ($p$ = .002 cluster-level FWE), left BNST ($p$ = .005 peak-voxel FWE-SVC), and right ventral striatum ($p$ = .005 peak-voxel FWE-SVC). In these areas, the approach-vs-avoid effect on BOLD was larger for passive compared to active responses, potentially signaling modulation of decision-related neural activity by motor preparation (**Supplementary Table S2**).

*Neural effects reflect decision (and not response) related changes in brain activity*

To verify whether our neural effects (main-text **Figure 3**) reflect decision-related rather than response-related changes in brain activation, we performed a finite impulse response (FIR) analysis to inspect the time course of the BOLD signal without assumptions on the shape of the hemodynamic response function (HRF). We focused on three ROIs that showed an effect for the voxel-wise approach vs. avoid contrast described above: the bilateral amygdala (AMY), ventral striatum (vStr), and vmPFC (**Supplementary Figure S1**). Three-way repeated measures ANOVAs revealed significant main effects of time and choice on BOLD in the amygdala and ventral striatum ($time_{AMY}$: F(1.1045, 62.9558) = 4.259, $p$ = .0137, $\eta_p^2$ = .069; $time_{vStr}$: F(0.9509, 54.2004) = 22.028, $p$ < .001, $\eta_p^2$ = .278; $choice_{AMY}$: F(0.36816, 20.9853) = 12.205, p < .001, $\eta_p^2$ = .176; $choice_{vStr}$: F(0.3168, 18.0668) = 8.6332, $p$ = .0046, $\eta_p^2$ = .132), a marginally significant main effect of choice in the vmPFC (F(0.31522, 17.96754) = 3.8532, $p$ = .054, $\eta_p^2$ = .063), and an interaction effect between time and choice for all three ROIs (AMY: F(1.1045, 62.9558) = 5.0339, $p$ = .003, $\eta_p^2$ = .081; vStr: F(0.9509, 54.2005) = 9.4897, $p$ < .001, $\eta_p^2$ = .142; vmPFC: F(0.94566, 53.90262) = 7.8983, $p$ < .001, $\eta_p^2$ = .121). More specifically, in all three regions the choice effect on BOLD was most pronounced in the bin spanning 4.5 – 6 seconds after stimulus onset (AMY: t(57) = 4.1693, $p$ < .001 ; vStr: t(57) = 4.6589; $p$ < .001; vmPFC: t(57) = 3.5415; $p$ < .001). In none of the regions there was a significant main effect of, nor an interaction effect involving, active vs. passive responses. These results further support the suggestion that the voxel-wise differences described above reflect value-based decision (and not response) related changes in BOLD response patterns (**Supplemental Figure S1**).

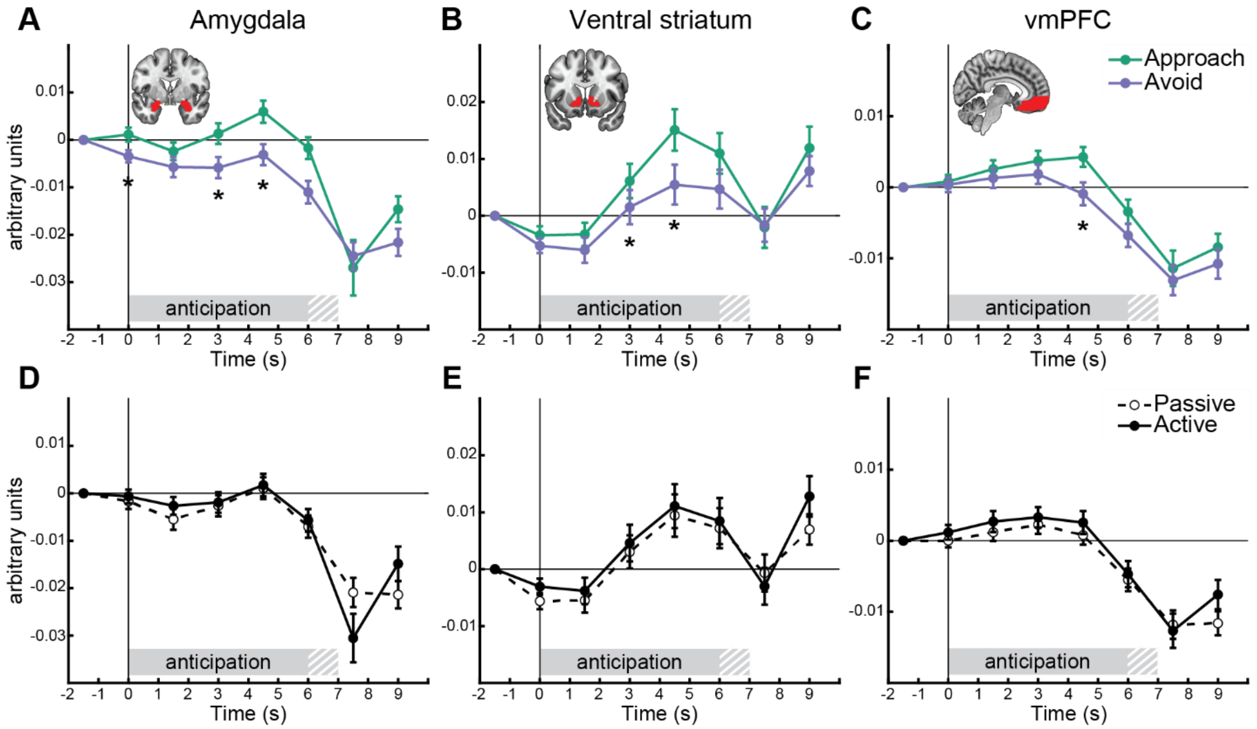


***Supplementary Figure S1. Time course trajectories of BOLD activity in the amygdala, ventral striatum, and vmPFC during the anticipation screen.*** *Stronger BOLD activity in approach compared to avoid trials in the amygdala (**A**), ventral striatum (**B**), and vmPFC (**C**) occur in time bins during the anticipation screen. Additionally, BOLD activity did not significantly differ as a function of subsequent active vs. passive responses (**D-F**). Each bin spans a time window of 1.5 s (e.g., bin 1 ranges from t = -1.5 to t = 0). Responses are plotted relative to the onset of the stimulus screen and baseline corrected relative to the first bin, indicating that neural effects are strongest during decision rather than the response. Error bars indicate +/- 1 SEM. Gray-white striped shaded areas reflect partial overlap between stimulus and target movement screens across different trials (i.e., movement window onset was uniformly jittered between 6 – 7 s). Asterisks indicate significant (p < .05) follow-up t-tests. vmPFC: ventromedial prefrontal cortex. Anatomical ROIs used to extract the BOLD signal are plotted in red.*

### *Formal comparison of freezing and base models*

We compared the fit of the three freezing models amongst themselves and with the base model. We used formal model comparison to find out to which of these models seems fits the choice behavior best. The comparison showed that while all models performed very similarly (i.e., the standard errors of the model comparison metrics between models overlapped), the *aversive value* model outperformed the base model as well as the other freezing models (i.e., the 'looic' was lowest, see **Supplementary Table S3**).

**Supplementary Table S3. Model comparison results of base and freezing models**

| **Model** | looic | $se_{looic}$ | $ELPD_{diff}$ | $se_{diff}$ |
|---|---|---|---|---|
| Base | 6049.105 | 110.4498 | -.04720 | 2.4694 |
| **Route 1 (AV)** | **6048.161** | **110.6306** | **0** | **0** |
| Route 2 (VC) | 6049.457 | 110.5524 | -0.6479 | 1.9312 |
| Route 3 (AI) | 6048.531 | 110.5737 | -0.1852 | 3.2587 |

*Lower model fit estimates (looic) indicate better fit. ELPD is the theoretical expected log pointwise predictive density for a new data set, estimated through leave-one-out (loo) cross validation. These estimates are compared between models using the $ELPD_{diff}$ metric, which reflects each model's ELPD relative to the best fitting model (in this case AV; more negative $ELPD_{diff}$ values indicate worse fit). The model with the best model fit is highlighted in bold.*

**Supplementary Table S4. Coordinates and statistics of positive and negative correlations between BOLD and model-based approach-avoidance during the anticipation screen**

| Region | Side | x, mm | y, mm | z, mm | Cluster size, $mm^3$ | Peak T | *p* value |
|---|---|---|---|---|---|---|---|
| *Base model – positive* | | | | | | | |
| Occipital cortex | L | -18 | -98 | 0 | 8,328 | - | < .001 |
| | R | 4 | -82 | -10 | 2,720 | - | < .001 |
| vmPFC | L/R | 4 | 64 | -2 | 4,416 | - | < .001 |
| Ventral striatum | L | -8 | 10 | -8 | - | 4.96 | = .001* |
| | R | 6 | 10 | -4 | - | 4.23 | = .009* |
| | R | 10 | 10 | -8 | - | 3.97 | = .019* |
| Amygdala | R | 32 | -2 | -18 | - | 3.87 | = .019* |
| *Base model – negative* | | | | | | | |
| Precuneus | R | 8 | -52 | 52 | 2,136 | - | <.001 |
| Superior frontal gyrus | R | 20 | 4 | 62 | 1,448 | - | = .008 |
| *AV – negative* | | | | | | | |
| Middle temporal cortex | R | 56 | 4 | -30 | 1,208 | - | = .007 |
| Paracentral lobule | R | 6 | -30 | 72 | 2,432 | - | < .001 |
| Inferior frontal cortex | L | -56 | 14 | 26 | 1,344 | - | = .004 |
| dlPFC | L | -34 | 4 | 62 | 1,128 | - | = .010 |
| Amygdala | L | -16 | -2 | -12 | - | 3.33 | = .086* |
| | R | 34 | 2 | -22 | - | 3.87 | = .022* |
| | R | 26 | 0 | -14 | - | 3.36 | = .081* |
| | R | 30 | -2 | -26 | - | 3.28 | = .096* |
| *AV – negative-avoidance* | | | | | | | |
| Amygdala | L | -2230 | 2 | -16 | - | 3.58 | = .045* |
| | R | 30 | -2 | -26 | - | 4.02 | = .014* |
| | R | 26 | 2 | -22 | - | 3.92 | = .018* |
| | R | 26 | 0 | -12 | - | 3.77 | = .027* |
| *VC – positive* | | | | | | | |
| dmPFC (dACC/SMA) | L/R | 2 | -6 | 70 | 17,664 | - | < .001 |
| Postcentral gyrus | L | -48 | -12 | 28 | 3,064 | - | < .001 |
| Middle occipital cortex | L | -24 | -72 | 24 | 1,640 | - | = .002 |
| Inferior temporal cortex | R | 52 | -10 | 26 | 2,016 | - | < .001 |
| Caudate nucleus | R | -16 | -2 | 22 | 1,008 | - | = .025 |
| Cerebellum | L | -42 | -56 | -24 | 2,256 | - | < .001 |
| Fusiform gyrus | R | 40 | -60 | -18 | 688 | - | = .035 |
| *VC – positive-approach* | | | | | | | |
| dmPFC (dACC/SMA) | L/R | -8 | 14 | 34 | 1,472 | - | = .004 |
| dACC | L/R | -8 | 14 | 34 | - | 4.93 | < .001* |

*All coordinates are defined in MNI152 space. All listed statistics are significant at $p < .05$ FWE-corrected at the cluster-level (whole-brain) or peak-level small volume corrected (SVC; predefined ROIs, indicated* *with asterisks '*'). For ROIs only, peak voxels with .05 < $p_{FWE}$ < .1 are also listed. dmPFC: dorsomedial prefrontal cortex; dACC: dorsal anterior cingulate cortex; SMA: supplemental motor area; vmPFC: ventromedial prefrontal cortex; dlPFC: dorsolateral prefrontal cortex.*
